# Proteus: Exploring Protein Structure Generation for Enhanced Designability and Efficiency

**DOI:** 10.1101/2024.02.10.579791

**Authors:** Chentong Wang, Yannan Qu, Zhangzhi Peng, Yukai Wang, Hongli Zhu, Dachuan Chen, Longxing Cao

## Abstract

Diffusion-based generative models have been successfully employed to create proteins with novel structures and functions. However, the construction of such models typically depends on large, pre-trained structure prediction networks, like RFdiffusion. In contrast, alternative models that are trained from scratch, such as FrameDiff, still fall short in performance. In this context, we introduce Proteus, an innovative deep diffusion network that incorporates graph-based triangle methods and a multi-track interaction network, eliminating the dependency on structure prediction pre-training with superior efficiency. We have validated our model’s performance on *de novo* protein backbone generation through comprehensive in silico evaluations and experimental characterizations, which demonstrate a remarkable success rate. These promising results underscore Proteus’s ability to generate highly designable protein backbones efficiently. This capability, achieved without reliance on pre-training techniques, has the potential to significantly advance the field of protein design. Codes are available at https://github.com/Wangchentong/Proteus.

## 1. Introduction

The biological function of a protein is often directly determined by its tertiary structure, which underscores the importance of designing novel protein backbones. *De novo* protein design methods are dedicated to creating proteins with the desired structure and function. Recent advancements in protein structure prediction methods, such as AlphaFold2 (Jumper et al., 2021) and RosettaFold (Baek et al., 2021), have enabled the ‘hallucination’ approach (Anishchenko et al., 2021; Wang et al., 2022) to directly generate protein sequences using backpropagation on the structure prediction networks. Further leveraging the generative capabilities of the diffusion model (Ho et al., 2020), RFdiffusion (Watson et al., 2023) has demonstrated superior performance across a wide range of protein design challenges. These applications include designing protein binders, scaffolding motifs, and creating symmetric oligomers. Despite RFdiffusion’s impressive performance, its dependency on pretraining with RosettaFold2 (Baek et al., 2023) poses a challenge for dissecting and refining the model’s architecture to improve performance for structure generation tasks.

To tackle the challenge of generating designable protein backbones without reliance on pretraining, researchers have developed a range of diffusion strategies and model architectures. One such approach is FoldingDiff (Wu et al., 2022), which employs diffusion in the protein backbone torsion space with a bidirectional transformer. This network iteratively denoises a sequence of torsion angles to generate protein-like backbones. However, the majority of generated structures are predicted to be non-designable. In contrast, two concurrent studies have shown more promise by directly applying diffusion on residue coordinates (Lin & AlQuraishi, 2023) or tangent space of coordinate and rotation (Yim et al., 2023). Additionally, previous research (Anand & Achim, 2022; Lee et al., 2023; Trippe et al., 2022) has explored multiple network architectures, including UNet (Ronneberger et al., 2015), Equivariant Graph Neural Networks (EGNNs) (Satorras et al., 2022), Invariant Point Attention(IPA) (Jumper et al., 2021), which have achieved success in diverse fields such as Computer Vision or dynamic system modeling.

Although these initiatives have moderately improved the designability of protein structure diffusion models without dependency on pretraining, there is still a notable performance gap compared to RFdiffusion, which results in considerable limitations for these models, making them less effective or difficult to apply in practical protein design tasks. RFdiffusion, with its cutting-edge capabilities, sets a higher standard in the field and highlights the need for further advancements in model development.

**Figure 1:**
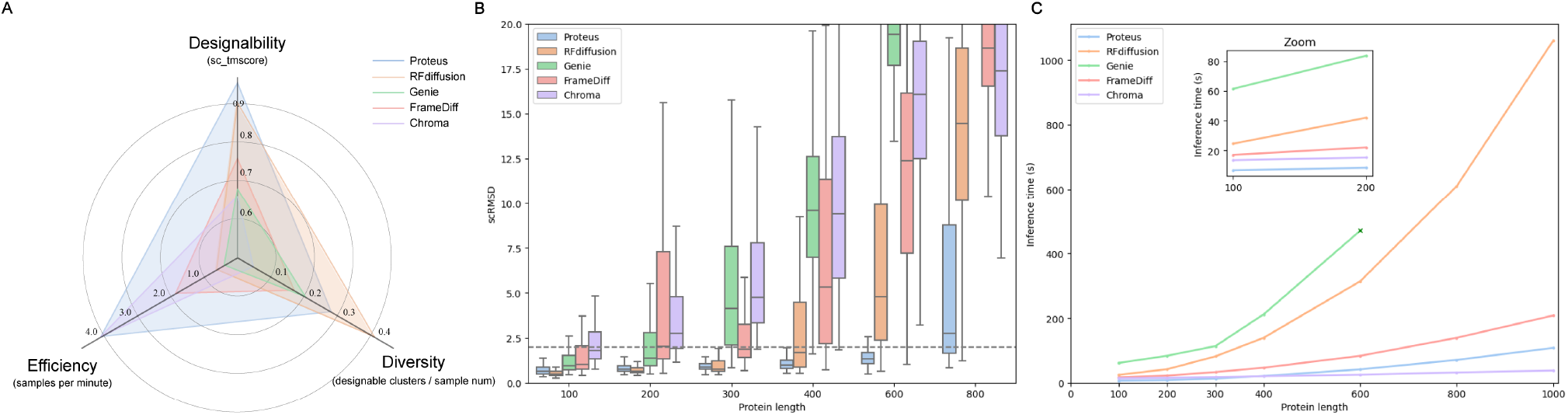
Benchmarking Proteus against other backbone diffusion models on designability, efficiency, and diversity. All metrics are averaged from 200 backbones of each length 100, 200, 300, 400, 600, and 800. For each backbone length, 8 sequences are designed by ProteinMPNN, except for Chroma, which uses ChromaDesign to achieve the best performance reported in its original paper. (A) Radar plot illustrating model evaluation across three dimensions. (B) Self-consistency RMSD between the generated backbone and the best prediction of ESMfold. (C) Sampling time of backbone generation, evaluated on A40. Genie fails to generate backbones larger than 600 residues due to running out of memory.

To bridge the performance gap between models that do and do not require pretraining, we have developed Proteus. Proteus achieves backbone designability on par with RFdiffusion by integrating a graph-based triangle technique and a multi-track interaction network, significantly bolstered by data augmentation. Moreover, our model sets a new efficiency standard, owing to two principal advancements: a reduction in the required sampling steps due to the enhanced representational capacity of the model, and the employment of local graph modeling to decrease computational complexity. These innovations allow Proteus to achieve protein generation speeds comparable to those of Chroma (Ingraham et al., 2022).

In summary, our model achieves several optimization objectives critical to protein design. It functions independently of pretraining, delivers high designability of protein structures, and maintains rapid generation speeds. The introduction of Proteus represents a noteworthy progression in protein design methodology, offering a solution that balances efficiency with the complexity of protein structure generation.

## 2. Preliminaries

### Protein backbone representation

Following the approach of AlphaFold2 (Jumper et al., 2021), the backbone of each residue is parameterized as a series of rigid transformations, also known as frames. These frames, denoted by T = (*R, t*) are defined within the special Euclidean group *SE*(3) and represent orientation-preserving transformations of the idealized backbone atom coordinates 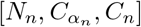. Specifically, *R* ∈ *SO*(3) is a rotation matrix derived from the backbone atoms N, CA, and C through the Gram-Schmidt process, and *t ∈* ℝ^3^ represents the coordinates of atom *C*_α_.

### Diffusion modeling on protein backbone

Multiple methodologies are available for protein backbone diffusion, including diffusion on inter-residue geometry or backbone torsion angles. In this paper, we detail our employed approach: SE(3) diffusion. This method treats each residue independently and computes the associated probability estimation on SO(3) for the score matching calculation, as originally proposed by Yim et al..

Briefly, the protein backbone’s forward diffusion process is driven by Brownian motion on SO(3) and ℝ ^3^ individually as shown in Equation (1)

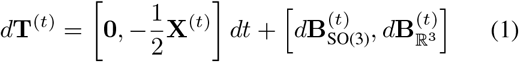

By defining probability estimation on ℝ^3^ forward diffusion, *p*_*t*|0_(*x*^(*t*)^|*x*^(0)^) = 𝒩 (*x*^(*t*)^; *e*^*−t/*2^*x*^(0)^, (1*−e*^*−t*^)*Id*_3_), the corresponding conditional score can be computed explicitly

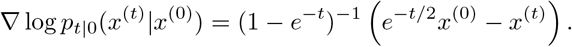

For the proper estimation of probability on SO(3), Brownian motion on SO(3) is defined as 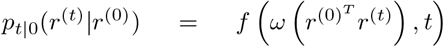 where ω(*r*) is the rotation angle in radians for any r *∈ SO*(3).The final probability estimation can be described as

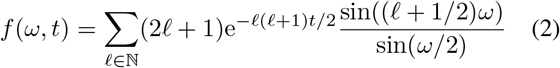

**Figure 2:**
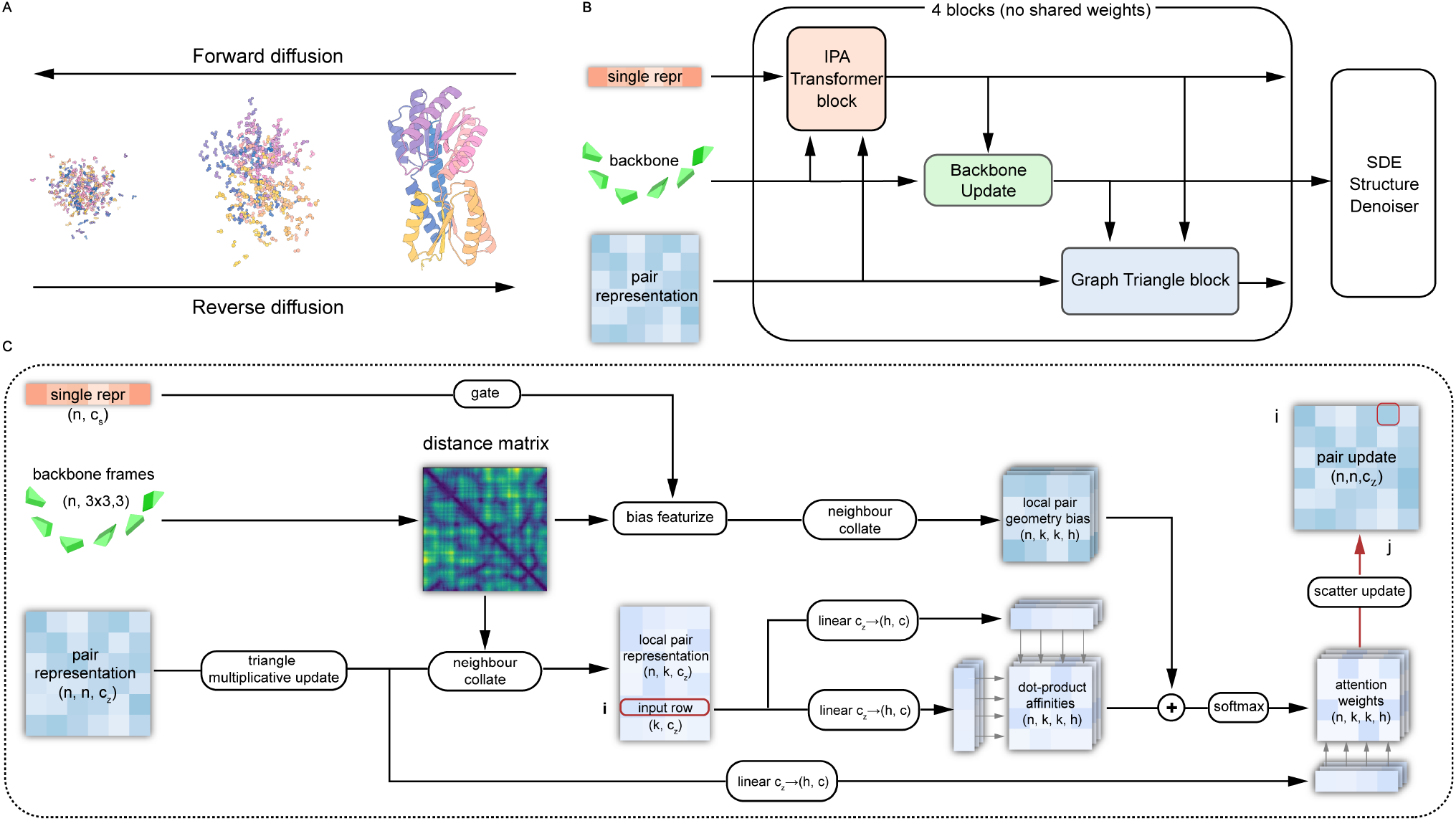
Model illustration (A) The Protein backbone diffusion model is trained to recover noised structures and generate new structures by reversing the forward process. (B) The overall architecture of Proteus. (C) The detailed model architecture of graph triangle block.

With the probability estimation, the corresponding conditional score can be computed as

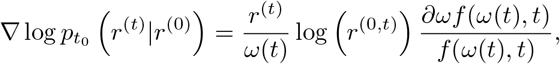

With a well-trained denoising score matching(DSM) network on tangent space of ℝ^3^ and SO(3), a protein backbone can be sampled from noise with n steps by Euler–Maruyama discretization (Bortoli et al., 2022).

In summary, the objective of the network is to predict a denoised protein backbone at timestep 0, given noisy backbones from any timesteps. Within this framework, probability estimation can be calculated, enabling the application of the stochastic differential equation (SDE) to sample a denoised protein backbone. We recommend that readers check Yim et al.‘s work for a profound understanding of SE(3) diffusion.

### Deep learning network architectures for protein structure modeling

The model architectures of AlphaFold2 (Jumper et al., 2021) and Rosettafold (Baek et al., 2021) have significantly advanced the development of protein backbone generation models. To elucidate the inspiration for our work, we provide a detailed account of these networks. AlphaFold2 utilizes the Evoformer module to process multiple sequence alignment (MSA) and structure template information into sequence and edge representations. Then, the Structure module iteratively refines these latent representations into the final protein structure, beginning from an initial ‘black hole’ configuration. The iterative nature of the Structure module, where a structure is input and refined through cycles, makes it a suitable backbone generation network for protein backbone diffusion (Anand & Achim, 2022; Yim et al., 2023; Lin & AlQuraishi, 2023). Similarly, the SE(3)-transformer (Fuchs et al., 2020) adopted by RosettaFold shares the iterative refinement capability. In addition to the iterative update mechanism, AlphaFold2’s heightened prediction precision is largely attributed to its triangle attention layer. This transformer-like architecture updates the edge representation of residue pairs by integrating the information of the third edge. However, due to its O(*N* ^3^) computational complexity, most current protein backbone diffusion models adopt a standard message passing layer or Unet (Ronneberger et al., 2015) for updating the edge representations. An exception is RFdiffusion (Watson et al., 2023), which employs axial attention (Ho et al., 2019) on residue pairs, thereby enhancing its representational capacity for the protein backbone denoising task.

## 3. Methods

### Algorithm 1 Proteus Model Inference

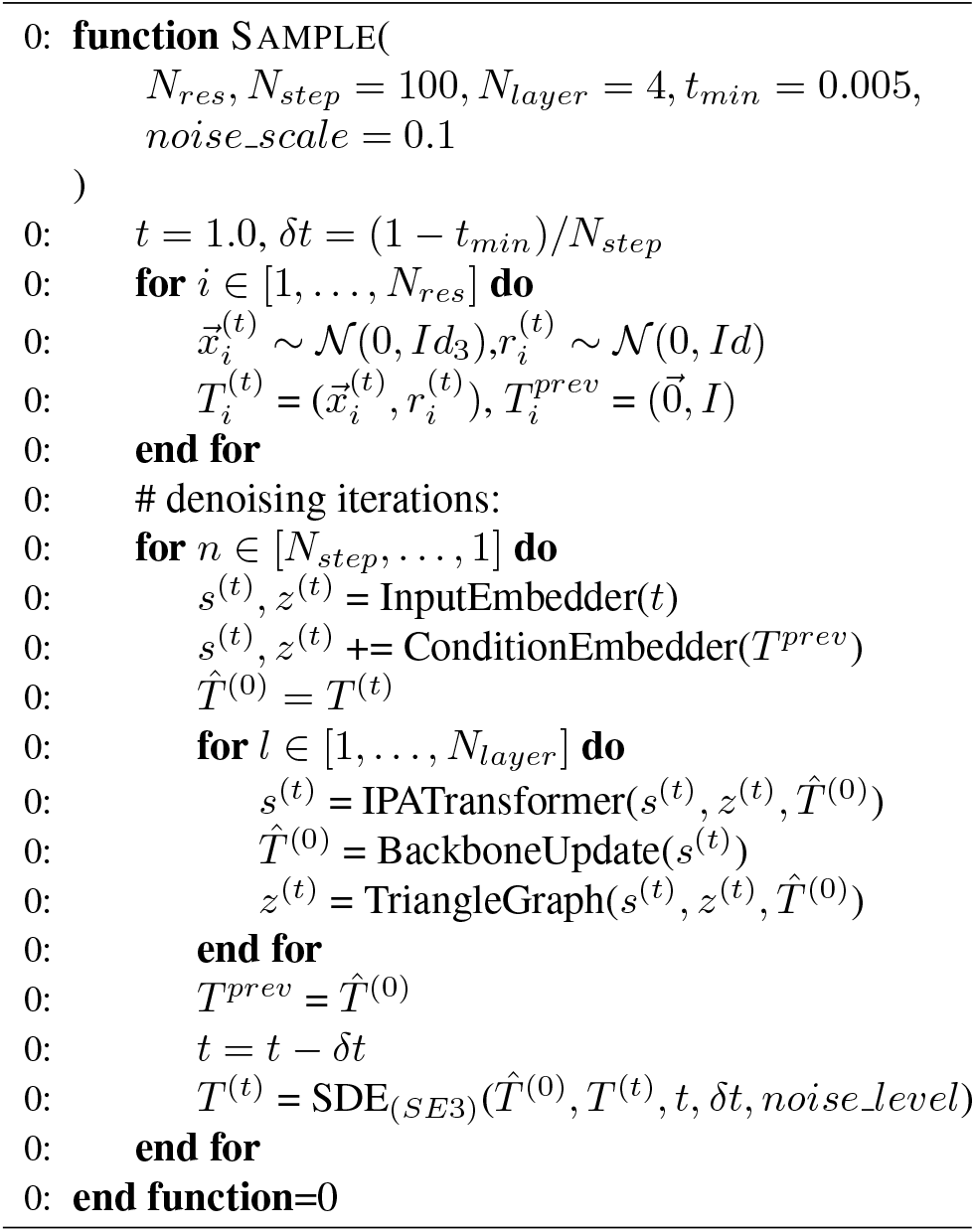

### 3.1. Model Architecture

#### Overview

In this section, we present the architecture of Proteus. Proteus iteratively updates the structural frames of proteins through a sequence of *L* layers of folding blocks. As shown in Figure 2B, each layer of the folding block receives input from three distinct tracks: node representation, edge representation, and structural frames. A folding block is composed of three components: an IPA-Transformer block, a backbone update layer, and a graph triangle block. Each component is tailored to model and refine one of input tracks while being aware of the representations from other tracks.

The IPA-Transformer block integrates an Invariant Point Attention (IPA) mechanism with a traditional transformer. The IPA conducts standard attention operations, incorporating a bias derived from the spatial distance between inter-residue atoms and edge representation. The Backbone Update, inspired by AlphaFold2’s methodology, utilizes a linear layer to predict translation and rotation updates for the frames of each residue, informed by the updated sequence representation.

The graph triangle block is tasked to update the edge representation. It employs a graph-based attention mechanism that operates on edge representation, modulated by a sequence representation-gated structural bias. The entire network consists of *L* layers of folding blocks, and no weights are shared among them. The sequence representation is initialized using the diffusion timestep and the edge representation is initialized using a relative sequential distance map introduced in AlphaFold2. When self-conditioning data is available, additional features, including the Ca distance map and relative rotational features from preceding predictions, are incorporated, with more elaborate explanations provided in Table 5. Our primary emphasis is on elucidating the graph triangle block, which is the source of significant enhancements in designability and efficiency.

#### Graph triangle block

The graph triangle block in our model is engineered to update the edge representation in the backbone diffusion process. Drawing inspiration from AlphaFold2’s network, we have innovatively adapted the concept of triangle attention and the multiplication method, which are central to AlphaFold2’s Evoformer module. Triangle attention is crucial for generating valid protein structures by enforcing the triangle inequality for the pair distance distribution when updating edge representations. This module significantly contributes to AlphaFold2’s exceptional accuracy but presents computational challenges due to its *O*(*N*^3^) computational complexity and substantial memory demands. Processing a 384-residue protein, a single layer of Evoformer can require upwards of 20GB memory during training, making it necessary to employ techniques of gradient checkpoint, especially considering the network comprises 48 Evoformer blocks. In terms of diffusion modeling, generating a 384 residue protein with 50 steps using AlphaFold2 can take a minimum of 10 minutes on a V100 GPU. Another challenge encountered with the application of the naive triangle attention technique to protein backbone diffusion models is the lack of awareness of the current structure. The Evoformer is tailored to convert Multiple Sequence Alignment (MSA) data into individual and edge representations, yet it lacks provision for structural information as an input to inform these update processes.

To address the aforementioned two major limitations when applying the triangle technique to protein backbone diffusion, we have devised the graph triangle block. This block computes triangle attention for edge representation with an optimized *O*(*NK*^2^) complexity. Utilizing the noisy input structure, we identify the k nearest neighbor residues for each residue and subsequently gather the *N ∗ K* edges from the comprehensive *N* ^2^ edge representation. Attention logits are then calculated among each residue’s *K* edges. To incorporate 3D spatial information, we calculate the inter-atom distances of the third edge and derive their Radial Basis Function (RBF) features as structural bias, rather than directly utilizing the representation of the third edge. Furthermore, the structure bias is gated by a feedforward network that leverages the sequence representations from both the starting and ending residues, thus ensuring seamless integration of inputs across all three tracks. Before triangle attention is performed for *N ∗ K* edge representations, we first apply a triangle multiplicative update, as used in Evoformer, to update the entire *N ∗ N* edge representations.

Employing this technique confers three distinct benefits over the axial attention layer used in RFdiffusion and the simpler message-passing layer utilized in FrameDiff and Genie. Firstly, the integration of triangle attention for edge representation updates markedly augments the model’s proficiency in modeling the protein backbone diffusion process. This advancement enables the model to achieve superior results with significantly fewer steps, yielding a threefold speedup and significantly outperforms the baseline model. Secondly, this approach significantly curtails memory requirements, allowing for training on much larger proteins without the necessity to crop the input. We can handle proteins up to 1024 residues in length during training on A40 GPU, compared to the 384 residues-limit for AlphaFold2 and RFdiffusion. This capability enhances the model’s efficacy in generating larger protein structures. Lastly, by incorporating inputs from all three tracks to update edge representation and protein structure, the protein structure is refined in a holistic manner. The architecture is more compact and best suited for structure-to-structure tasks, such as protein structure generation and predicting the protein’s apo-to-holo conformational transition (Hou et al., 2023).

Employing this technique confers three distinct benefits over the axial attention layer used in RFdiffusion and the simpler message-passing layer utilized in FrameDiff and Genie. Firstly, integrating triangle attention for edge representation updates significantly enhances the model’s proficiency in modeling the protein backbone diffusion process. This advancement enables the model to achieve superior results with significantly fewer steps, yielding a threefold speedup and markedly outperforming the baseline model. Secondly, this approach substantially reduces memory requirements, allowing for training on much larger proteins without the necessity of cropping the input. We can handle proteins up to 1024 residues in length during training on A40 GPU, compared to the 384-residue limit for AlphaFold2 and RFdiffusion. This capability markedly augments the model’s efficacy in generating larger protein structures. Lastly, by incorporating inputs from all three tracks to update the edge representation and protein structure, the protein structure is refined in a holistic manner. The architecture is more compact and is ideally suited for structure-to-structure tasks such as protein structure generation. It also holds the potential to generalize to the protein’s apo-to-holo conformational transitions (Hou et al., 2023).

### 3.2 Training

#### Dataset

We curated a dataset from the Protein Data Bank (PDB) (Berman et al., 2000) with a cutoff date of August 1, 2023. Instead of the conventional practice of training diffusion models solely on monomeric structures, we also included oligomeric structures and extracted their individual chains as training data. To avoid redundancy, we mapped protein sequences to UniProt IDs and selected the highest resolution structure for chains that shared the same UniProt ID and exhibited at least 80 percent sequence overlap. Subsequently, we filtered out the remaining protein chains with length cutoffs between 60 and 512. Additionally, we limited the inclusion of proteins that contained a maximum of 50 percent loop regions as assigned by the DSSP program (Kabsch & Sander, 1983). This curation process yielded a training set of 50,773 single-chain proteins. Our results indicate that augmenting the dataset with additional singlechain data derived from oligomeric structures significantly enhances model performance, compared with training solely on monomers. Supporting evidence of this enhancement is presented in Table 4.

#### Training losses

We have adopted the training loss used in FrameDiff, which can be divided into two main components: denoising translation and rotation score-matching losses, and the auxiliary losses involving the pair-wise distance matrix and coordinate loss on backbone atoms, as depicted in Equation 3. The positions of oxygen atoms are calculated based on the coordinates of other backbone atoms.

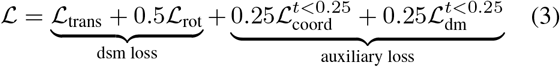

Here, ℒtrans computes the L2 loss between predicted translations and native translations, while ℒrot calculates the L2 loss on rotation scores weighted by 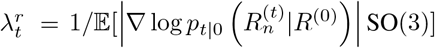, following the approach in (Song et al., 2020). ℒcoord represents the coordinate loss between predicted backbones and native backbones. Lastly, ℒ_dm_ computes the pair-wise distance loss between predicted atom positions and native positions. We apply auxiliary losses when t < 0.25. Empirically, we have observed that including auxiliary losses at a small timestep can aid convergence and enhance the model’s performance, which aligns with the findings of FrameDiff.

### 3.3 Sampling

The sampling procedure is detailed in Algorithm 1. We initiate the process by sampling the initial frames for rotation and translation separately. For the initial translation, we employ a Gaussian distribution on ℝ^3^. As for the rotation component, we first sample the coefficients of orthonormal basis vectors of the Lie algebra `so`(3) and translate them into the rotation matrix. Once initialized, Proteus takes the input noisy structure and generates a prediction structure at the timestep 0. Then, we iteratively apply the Euler–Maruyama discretization (Bortoli et al., 2022) as the SDE solver for *N*_steps_ to generate the denoised structure.

## 4. Experiments

We rigorously evaluated Proteus performance through both in-silico validation and in vitro experimental approaches. In Section 4.1, we show Proteus’ performance in the task of unconditional monomer generation. For a comprehensive assessment, we benchmark Proteus against a suite of leading protein backbone diffusion models, including Chroma (Ingraham et al., 2022), RFdiffusion (Watson et al., 2023), FrameDiff (Yim et al., 2023), and Genie (Lin & AlQuraishi, 2023). Furthermore, we extended our evaluation to the generation of protein complexes, where we compare Proteus’s efficacy in generating oligomers of dimer, trimer, and tetramer against Chroma.

Section 4.2 is dedicated to the in vitro experimental validation of Proteus. We synthesized the DNA oligos for 16 designed proteins generated by Proteus, expressed and characterized their biochemical properties, particularly folding and stability. The objective of these experiments is to substantiate the model’s practical utility and its effectiveness in real-world biological applications.

### 4.1. in-silico protein generation and evaluation

#### Monomer backbone generation and evaluation

To comprehensively evaluate the performance of a protein backbone diffusion model, it is essential to consider three primary factors: designability, efficiency, and diversity. These aspects are critical to the overall assessment and are detailed in Table 1

**Table 1:** Performance comparison of models on the unconditional monomer generation task. Designability is assessed by the self-consistency TM-score between generated and refolded backbones. Efficiency is quantified by number of seconds required to generate a sample, while diversity is measured by the ratio of designable clusters to the total number of generated backbones. Results are averaged across backbone lengths of 100, 200, 300, 400, and 600, as detailed in Figure 1. The top-performing metrics are highlighted in bold, with the second-best results underlined.

| Method | $N_{\text{params}}$ | Designability (↑) | Sampling time (s) (↓) | Diversity (↑) | Timesteps (↓) |
| --- | --- | --- | --- | --- | --- |
| Proteus | 19.8M | **0.921** | **18.20** | <u>0.235</u> | 100 |
| RFDiffusion | 59.8M | <u>0.705</u> | 120.24 | **0.328** | 50 |
| GENIE(SwissProt) | 4.1M | 0.349 | 188.07 | 0.163 | 1000 |
| FrameDiff | 17.4M | 0.405 | 40.47 | 0.136 | 500 |
| Chroma | 18.5M | 0.174 | <u>18.31</u> | 0.038 | 500 |

**Designability** aims to measure the quality of the generated backbones, by designing sequences for the generated backbones and refolding them to compute the errors between the generated and folded structures. Designability is the foremost factor, indicating the likelihood of identifying a protein sequence to fold into the designated structure. This is the cornerstone metric for gauging the performance of a diffusion model, as it directly correlates to the model’s capacity to generate viable proteins that could conceivably exist in nature.

In our implementation, we use ProteinMPNN (Dauparas et al., 2022) at sampling temperature 0.1 to generate 8 sequences for the designed backbone. Specifically, for Chroma derived backbones, we employ its dedicated inverse folding model, ChromaDesign (Ingraham et al., 2022) at temperature 0.1 and diffusion augmentation 0.5. This substitution is made following the observation that ChromaDesign yields a higher success rate for Chroma’s backbones, as documented in its paper. First, the inverse folding model generates multiple sequences corresponding to the sampled backbone, which are subsequently fed into ESMFold (Lin et al., 2023) to fold the structure. The designability of the backbone is represented by the highest TM-score (scTM, evaluating the similarity between two structures, where higher values are better) in all the predicted structures. Additionally, we also compute the self-consistency C_α_-RMSD, where the threshold of < 2Å is the criterion for successful design. Any generated protein exceeding this C_α_-RMSD threshold is deemed non-functional and likely to be unsuccessful in experimental validation.

**Efficiency** is the next critical factor. It is a key determinant in the success of design models, especially when computational resources are limited. A more efficient model can generate more samples in a limited time which increases the likelihood of generating potent candidates. Given the limitations on computation resources, efficiency becomes a pivotal criterion, on par with designability.

Efficiency is estimated as the time taken to generate a protein backbone on a standard NVIDIA Ampere Tesla A40 GPU with 48 GB of GPU memory. Additionally, we propose a **Time-for-Success Design (TSD)** metric, which computes the time consumed to generate a designable backbone. This metric underscores the crucial balance between the time required to generate successful designs and the efficiency of the model, making it highly relevant for real-world applications where both speed and effectiveness are critical.

**Figure 3:**
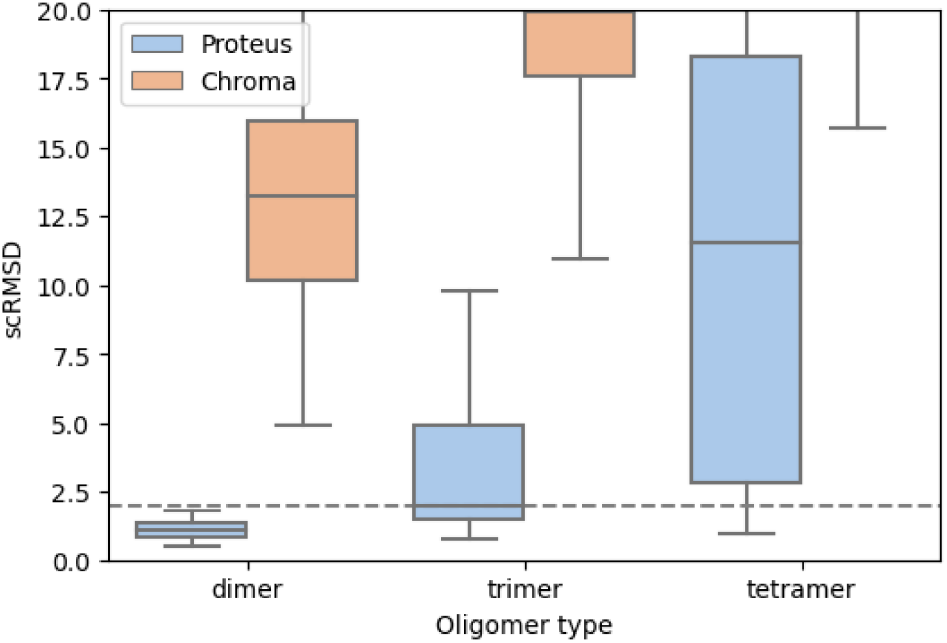
Complexes generation benchmark. scRMSD is derived from 200 backbone samples for each type of oligomer, with the length of each chain fixed at 200 amino acid residues.

**Diversity** is the third essential consideration, gauging the structural variance among the generated protein backbones. Diversity is evaluated by calculating the ratio of diverse **designable** structure clusters to the total number of generated backbones. Specifically, we cluster the generated backbones using MaxCluster (Herbert & Sternberg, 2008) with a TM-score cutoff of 0.6. Notably, we only consider backbones that are designable, defined as those with a C_*α*_RMSD of less than 2 Å. This criterion is important because sub-optimal models often produce unrealistic structures that cannot be folded by any sequence and are highly non-physical. Including such backbones in the assessment would skew the metric towards models generating random, sub-optimal structures rather than models that genuinely capture the realistic distribution.

#### Complexes backbone generation and evaluation

In Figure 3, we assess the capability of Proteus in generating protein complexes by drawing comparisons with Chroma under a set of monomer design evaluation metrics. We compute scRMSD across different oligomer configurations, including dimers (two chains), trimers (three chains), and tetramers (four chains), with each chain consisting of 200 amino acids. Notably, despite the model initially being trained on single protein chains, we observe its adaptive generation capacity to oligomers. This is achieved by adding a large positional index as a chain breaker (Watson et al., 2023), a technique elaborated in Appendix A.1. Additionally,Figure 6 offers visual representations of the oligomeric structures produced by this method.

#### Results

Proteus was benchmarked against the leading-edge protein backbone diffusion models, including RFDiffusion, Genie, FrameDiff, and Chroma. We generated 200 backbones across a spectrum of protein lengths, specifically [100, 200, 300, 400, 600], and computed the mean score for every category, as delineated in Table 1. Figure 1B shows the scRMSD distribution for each length category. Proteus matches the performance of RFdiffusion for shorter sequence lengths(100*−*300 residues) and significantly excels in generating sequences longer than 300 amino acids.

We attribute the enhanced capability to the novel architecture of Proteus, which employs graph-level triangle techniques. Furthermore, the enhanced performance of Proteus in oligomeric structure generation, surpassing that of Chroma, provides evidence of its robust out-of-distribution generative capabilities.

Remarkably, while upholding high designability, Proteus also matches Chroma in computational efficiency. Table 3 illustrates that Proteus, through enhanced network representation capabilities, requires merely 100 sampling steps without compromising its designability. This is in contrast to the 1,000 steps necessary for Genie, and 500 steps for FrameDiff and Chroma. Figure 1 showcases the computation time associated with varying protein lengths. Proteus exhibits faster performance compared to Chroma in generating proteins with lengths shorter than 400 amino acids. However, Chroma shows greater time efficiency on proteins exceeding 400 amino acids. Proteus, with superior designability and efficient graph-level processing, distinctively surpasses other models in the realm of success efficiency, as evidenced by the benchmark results in Table 2.

**Table 2:** Comparison of the Time-for-Success Design (TSD) Metric. This metric evaluates the time required to generate a designable backbone (scRMSD < 2Å) on an A40. Asterisks (*∗*) indicate models that failed to predict any designable backbones within 200 samples. The best results are bolded.

| Protein length | TSD (s) |  |  |  |  |  |
| --- | --- | --- | --- | --- | --- | --- |
|  | 100 | 200 | 300 | 400 | 600 | 800 |
| Proteus | **7.02** | **9.11** | **13.47** | **23.09** | **50.62** | **196.11** |
| RFDiffusion | 24.84 | 43.97 | 97.14 | 262.64 | 1494.28 | 30500.00 |
| GENIE (SwissProt) | 72.66 | 127.55 | 482.97 | 21169.0 | * | * |
| FrameDiff | 23.12 | 45.12 | 62.11 | 216.88 | 1851.11 | * |
| Chroma | 24.88 | 56.66 | 352.00 | 4020.00 | * | * |

**Table 3:** Proteus sample parameter benchmark

|  |  |  |  |  |  |  |
| --- | --- | --- | --- | --- | --- | --- |
| noise level $\zeta$ | 0.1 | 0.5 | 1.0 | 0.1 | 0.1 | 0.1 |
| $N_{\text{timestep}}$ | 100 | 100 | 100 | 50 | 200 | 500 |
| < 2Å scRMSD (↓) | 92.0% | 87.8% | 37.3% | 89.8% | 91.0% | 92.2% |
| DIVERSITY (↑) | 0.23 | 0.27 | 0.16 | 0.26 | 0.22 | 0.22 |

#### Ablation

In our comprehensive ablation analysis, as delineated in Table 4, the enhancements are attributed to three main aspects: dataset augmentation, self-condition enhancement, and model architecture. Previous protein backbone diffusion methods offered a range of training datasets: FoldingDiff utilized 30,395 protein chains from the CATH database (Sillitoe et al., 2014) with a cropping length of 128. Genie adopted the AlphaFold-Swissprot database (Varadi et al., 2021), containing 195,214 protein chains with a pLDDT cutoff of 80 and a length cutoff of 256. FrameDiff (Yim et al., 2023) compiled a dataset from PDB entries of only monomers, capped at 512 amino acids, resulting in a collection of 20,312 chains. In contrast to the above approaches, we expanded the dataset by adding oligomeric structures, which were subsequently split into single chains, resulting in a dataset of 50,773 chains that significantly bolstered designability. The ablation study further reveals the benefits of incorporating pairwise rotational info as a self-conditioning feature, analogous to the template featurization introduced by AlphaFold2, enabling the model to capture the structure self-consistency from previous denoising steps more accurately.

**Table 4:**
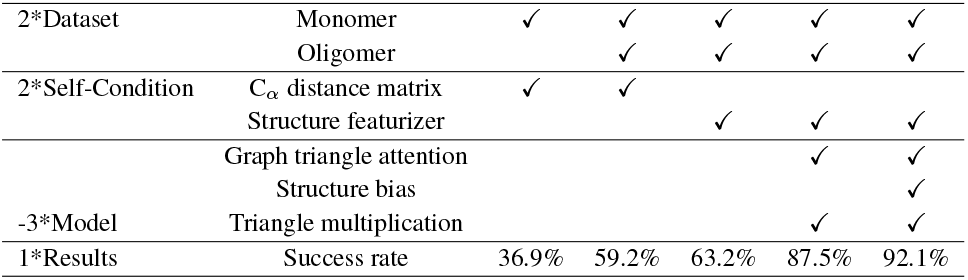
Ablation Study

While these incremental modifications substantially elevated the model’s performance, it was the introduction of the local triangle graph technique that allowed Proteus to match the performance of RFDiffusion. This innovative network, which applies an attention mechanism at the graph level with the integration of structure bias, not only preserves a fast sampling speed on par with Chroma but also significantly improves designability. This indicates that structure-level information can enrich edge representation, a novel insight not yet explored by AlphaFold2.

### 4.2 Experimental validation

DNA oligonucleotides encoding 16 designs generated by Proteus were synthesized, and the proteins were recombinantly expressed in Escherichia coli. This set comprises 12 proteins of 300 amino acids and 4 proteins of 500 amino acids. All proteins were well expressed in E. coli. Size exclusion chromatography (SEC) analysis revealed monodisperse peaks for 9 of the 300 amino acid designs and 3 of the 500 amino acid designs, which corresponded to the expected molecular weight. Furthermore, circular dichroism (CD) spectroscopy confirmed the well-folded structure of these designs, and the secondary structure features are consistent with the design models. Notably, these proteins exhibited remarkable thermostability, remaining well-folded at temperatures up to 95°C. Experimental results are comprehensively detailed in Figure 4.

## 5. Related work

### Structure diffusion models on proteins

Motivated by the significant achievements of diffusion models (Ho et al., 2020; Song et al., 2020; Bortoli et al., 2022) protein diffusion models have been developed to generate proteins in either sequence or structural space, with certain methods adeptly bridging both spaces (Anand & Achim, 2022). Anand & Achim pioneered a model that co-diffuses backbone, sequence, and sidechain information utilizing the Structure Module of AlphaFold2. Subsequently, numerous methods have been introduced, focusing on the diffusion of inter-residue geometry (Lee et al., 2023) and backbone dihedral angle (Wu et al., 2022). The leading edge protein diffusion models predominantly engage with diffusion in either SE3 or R3 space in an end-to-end fashion (Yim et al., 2023; Lin & AlQuraishi, 2023). These methods have been further expanded to function motif scaffolding (Trippe et al., 2022; Yim et al., 2024). Chroma achieved a higher efficiency by utilizing an efficient graph neural network. RFdiffusion (Watson et al., 2023) has attained state-of-the-art designability through finetuning the Rosettafold2 structure prediction network (Baek et al., 2023). In contrast to RFdiffusion’s approach, our work achieves a comparable designability and much higher efficiency without the necessity of pretraining, making a significant departure from existing methodologies.

**Figure 4:**
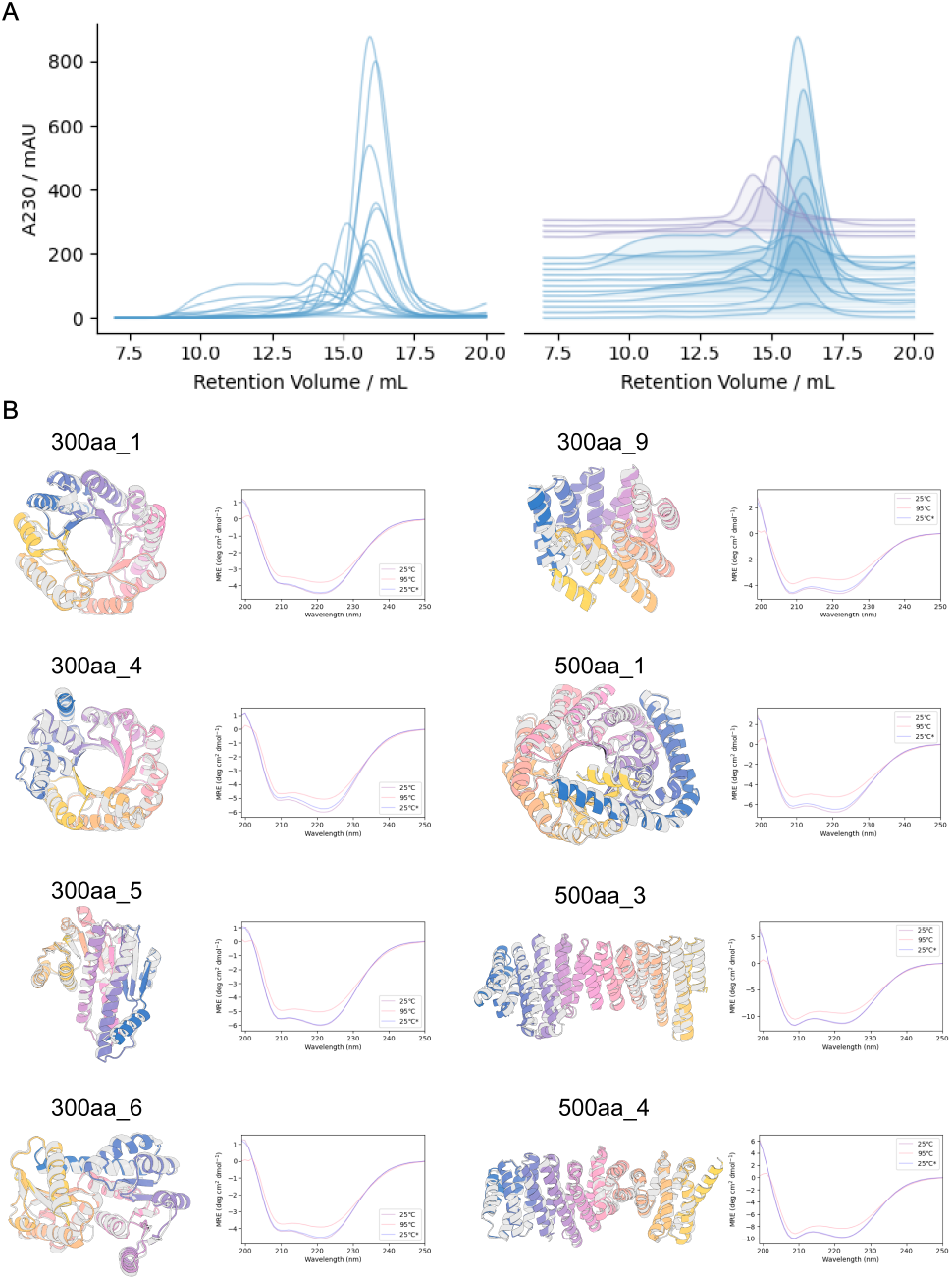
Experimental characterization of the generated proteins indicates they are well-folded monomers and thermostable. (A) Size exclusion chromatography profiles of the recombinantly expressed proteins on a Supderdex 75 Increase column. (B) Examples of designed proteins that are expressed as monomers. Designs (grey) overlaid with AF2 predictions (colors) are shown on the left, alongside circular dichroism (CD) spectra at 25°C, 95°C and 25°C (*) after cooling down from 95°C on the right.

### Other diffusion models on proteins

Many other studies have pivoted to the diffusion of protein sequences, or the synergistic integration of sequence data into structure diffusion models. EvoDiff (Alamdari et al., 2023), for example, utilizes evolutionary-scale protein sequence data to generate protein sequences. Although EvoDiff can generate quite diverse protein sequences, the structure prediction metrics indicate limited designability for the generated backbones. Chu et al. innovatively incorporates protein sequence features into the protein backbone diffusion process. The model utilizes a timestep-dependent ProteinMPNN model to design sequences on the noisy structures. However, this appears to diminish the model’s performance. This runs counter to the intuitive notion that the integration of sequence data should enhance structural quality. We notice another possible formulation of structure-sequence co-generation by language model inspired by SaProt (Su et al., 2023). For the specific task of antibody diffusion, considerable efforts (Luo et al., 2022; Kong et al., 2023; Peng et al., 2023) have been made to simultaneously generate the sequence and structure of complementarity-determining regions (CDR), highlighting the community interest in antibody design and the potential of sequence-structure co-diffusion.

### Protein-ligand complexes prediction

In previous works, the prediction of protein-ligand complexes is treated as a regression problem, focusing on the rigid body docking of ligands to holo proteins (Stärk et al., 2022; Lu et al., 2022). Capitalizing on the generative power of diffusion models, Diffdock significantly improved the docking accuracy (Corso et al., 2022). Protein-ligand co-folding models (Qiao et al., 2023; Lu et al., 2024) offer more precise predictions when the holo-state protein structures are not known a formidable and crucial challenge in drug discovery. Furthermore, RFdiffusion All Atom (Krishna et al., 2023) are trained to design novel small molecule binders, expanding the frontier of computational approaches in molecular design.

## 6. Discussion

In this paper, we introduce a new model architecture for protein backbone diffusion, and demonstrate its enhanced designability and efficiency without the necessity of pretraining. Our model advances the field by integrating the triangle attention technique into residue edge representation update and building multi-track interaction networks to enhance its representation capability. Our model shows improved performance in generating longer monomers (with 400 or more amino acids) compared to RFdiffusion. The successful generation of oligomeric structures further reveals Proteus’ generalizability.

Looking forward, we envision several research trajectories where Proteus could exert a transformative influence. Proteins in their natural state often manifest as multi-chain entities, orchestrating their functions in a coordinated fashion. Given Proteus’s exemplary performance in crafting larger proteins and multi-chain architectures with both high fidelity and efficiency, it is ideally suited for direct application in the generation of protein oligomeric structures and complex protein machinery. This capability will broaden the current scope of designable protein space, enabling the creation of innovative protein nanomachines.

A further domain of interest is the area of protein-ligand co-folding. Contemporary breakthroughs, exemplified by tools like Rosettafold All Atom (Krishna et al., 2023) and AlphaFold3 (Abramson et al., 2024), have eclipsed the previously established benchmarks set by Diffdock in the realm of modeling protein-small molecule interactions. The integration of Proteus-inspired methodologies into the diffusion dynamics of ligand-protein interplay holds the promise of refining protein-ligand co-folding techniques, paving the way for the development of methods with enhanced accuracy.

## Acknowledgements

The authors thank Pranam D. Chatterjee, Fajie Yuan, Jin Su, Jiaxuan Wang, all reviewers, the area chair for helpful discussions, Minchao Fang for his help with code development, and the Westlake University HPC Center for computing support.

This work was supported by the Science and Technology Innovation 2030 Major Project (2021ZD0150100), the National Key RD Program of China (2021YFC2301401, 2022YFA1303700), the National Natural Science Foundation of China (32370989), and the Westlake Education Foundation.

## Impact Statement

This paper presents work whose goal is to advance the field of protein design. There are many potential societal consequences of our work, none which we feel must be specifically highlighted here.

## A. Additional method details

Here, we describe the implementation details of Proteus as a further explanation of Section 3. To briefly revisit the fundamentals, the ℓ-th layer’s node representations are denoted as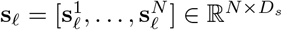, edge representations are captured by 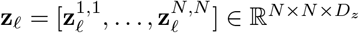. The spatial configuration of each residue in the ℓ-th layer is encapsulated by *T*_*ℓ*_ *∈* SE(3)^*N*^.

### A.1 Proteus model details

**Table 5:** Input feature of Proteus

| Input Feature (shape) | Description |
| --- | --- |
| aa ($N_{res}, 21$) | The amino acid type of input residues, include all 20 standard types with 1 unknown type |
| timestep ($N_{res}, 32$) | The timestep embedding from 0 to 1 by sinusoidal positional encoding. |
| rigid_frame ($N_{res}, 7$) | Input noisy structure, 3 dim of translation and 4 dim of rotation represented by a quaternion. |
| rel_pos ($N_{res}, N_{res}, 65$) | Relative position index embedding in [-32,32.] |
| torsion_angle_prev($N_{res}, 14$) | Self-condition input structure’s residue torsion angle of prev prediction represented by sin and cos formulation, 4 chi angles will be masked out in the network. |
| distogram_prev($N_{res}, N_{res}, 39$) | Self-condition input structure’s pair distance histogram of pseudo $C\beta$, bin_min=3.25Å, bin_max=50.75Å. |
| unit_vector_prev($N_{res}, N_{res}, 3$) | Self-condition input structure’s unit vector of each residue’s $C\alpha$ coordinate in another residue’s local frame. |

#### Feature initialization

Node feature is initialized from timestep and amino acid type (currently, all residues amino acid type is set to alanine for unconditional generation). The edge feature is initialized from the two corresponding node features with additional relative sequence position encoding. The Multi-Layer Perceptrons (MLP) used to embed initial features consists of 3 Linear layers with biases, 2 ReLU activation layers between Linear layers, and a LayerNorm (Ba et al., 2016) at the end.

The timestep embedding embedding ϕ(·) follows Ho et al. by using sinusoidal embeddings (Vaswani et al., 2023). The relative sequence position encoding *relpos*(·) follows algorithm 4 in supplementary of Alphafold2 (Jumper et al., 2021).

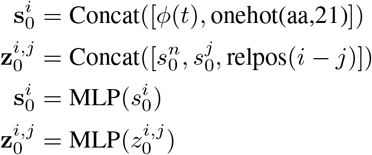

For the positional encoding, the relative position tokens of residue i and residue j within the same protein chain are always between -32, -31,…,31, 32. For oligomer generation, we add an extra position index 200 on the residues of the next chain, so the relative position tokens of residue i and residue j of different chains always have a position index -32 or 32, indicating a chain break.

#### Self-condition featurization

Encoding of predicted structure from the previous step has been proved to improve prediction self-consistency and backbone designability (Watson et al., 2023). In 50% time of training and all inference time, the ConditionEmbedder mentioned in Algorithm 1 is used to encode the prediction of the previous step as node embedding **t**_*s*_ and edge embedding **t**_*z*_, and the self condition feature is encoded to s_0_ and z_0_ after feature initialization.

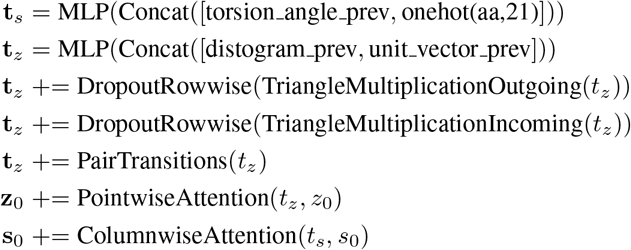

We have developed a self-conditioning featurization module, taking inspiration from AlphaFold2’s TemplatePairStack (Algorithm 16). The self-conditioned feature is divided into node features *t*_*s*_, which include predicted backbone dihedral angles and amino acid types, and edge features *t*_*z*_, which comprise *C*β pair distance and SE(3)-invariant pairwise directional vectors.

The node feature is encoded by an MLP and incorporated into s_0_ via ColumnwiseAttention. This architecture is similar to AlphaFold2’s MSAColumnAttention (Algorithm 8) but without gating. The edge feature is similarly encoded by an MLP and then processed through two triangle multiplication layers(AlphaFold2 Algorithm 11) with a dropout rate of 0.25 and a PairTransition layer (AlphaFold2 Algorithm 15). It is ultimately integrated into *z*_0_ through PointwiseAttention(Alphafold2 algorithm 17). Notably, we have reduced the number of blocks to one block and omitted the triangle attention layer present in AlphaFold2’s TemplatePairStack to enhance efficiency.

#### IPA-Transformer block

The IPA-Transformer block is used to update node information, and we will describe its details here. An IPA-Transformer block incorporates an Invariant Point Attention (IPA) as presented in AlphaFold2 (Jumper et al., 2021) without any alterations, along with a standard Transformer layer (Vaswani et al., 2023). IPA was first introduced by Anand & Achim as the central model architecture of the protein structure diffusion model. The Transformer is appended to IPA to enhance the representation ability proposed by Framediff (Yim et al., 2023).

The node update equations are as follows:

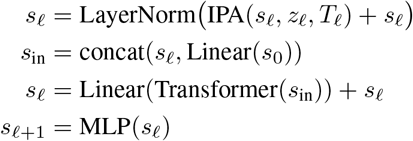

#### Backbone update

Our frame updates follow the BackboneUpdate algorithm in AF2(Algorithm 23). We write the algorithm here with our notation,

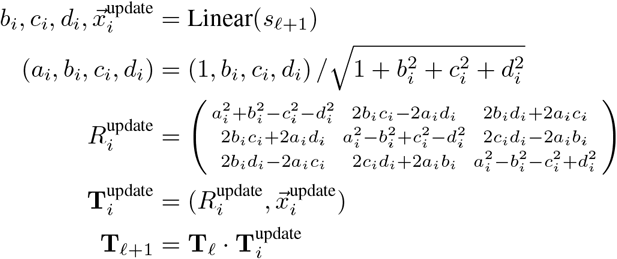

As shown in the equation, an unnormalized quaternion consists of *b*_*i*_, *c*_*i*_, *d*_*i*_ ∈ ℝ^3^ and a translation vector 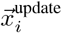 is predicted from s_*ℓ*+1_. We iterative update each residue’s frame by applying the translation 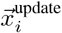 and rotation matrix 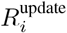 to **T**_*ℓ*_.

#### Graph triangle block

The graph triangle block is designed for the update of edge information. we further explain the implementation details of the graph triangle block mentioned in Section 3.1 and Figure 2. We develop the StructureBiasedGraphTriangleAttention layer(Algorithm 2,Algorithm 3) as the core network for local attention calculation with two triangle multiplication layers(AlphaFold2 Algorithm 11) for global update of edge feature.

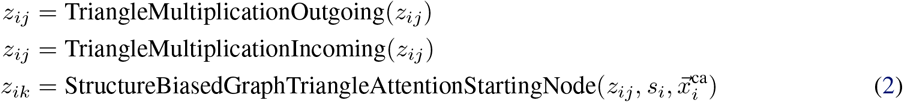

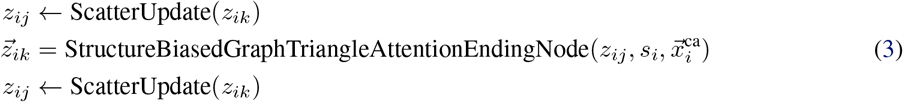

##### Algorithm 2

Structure biased graph triangle attention around starting node

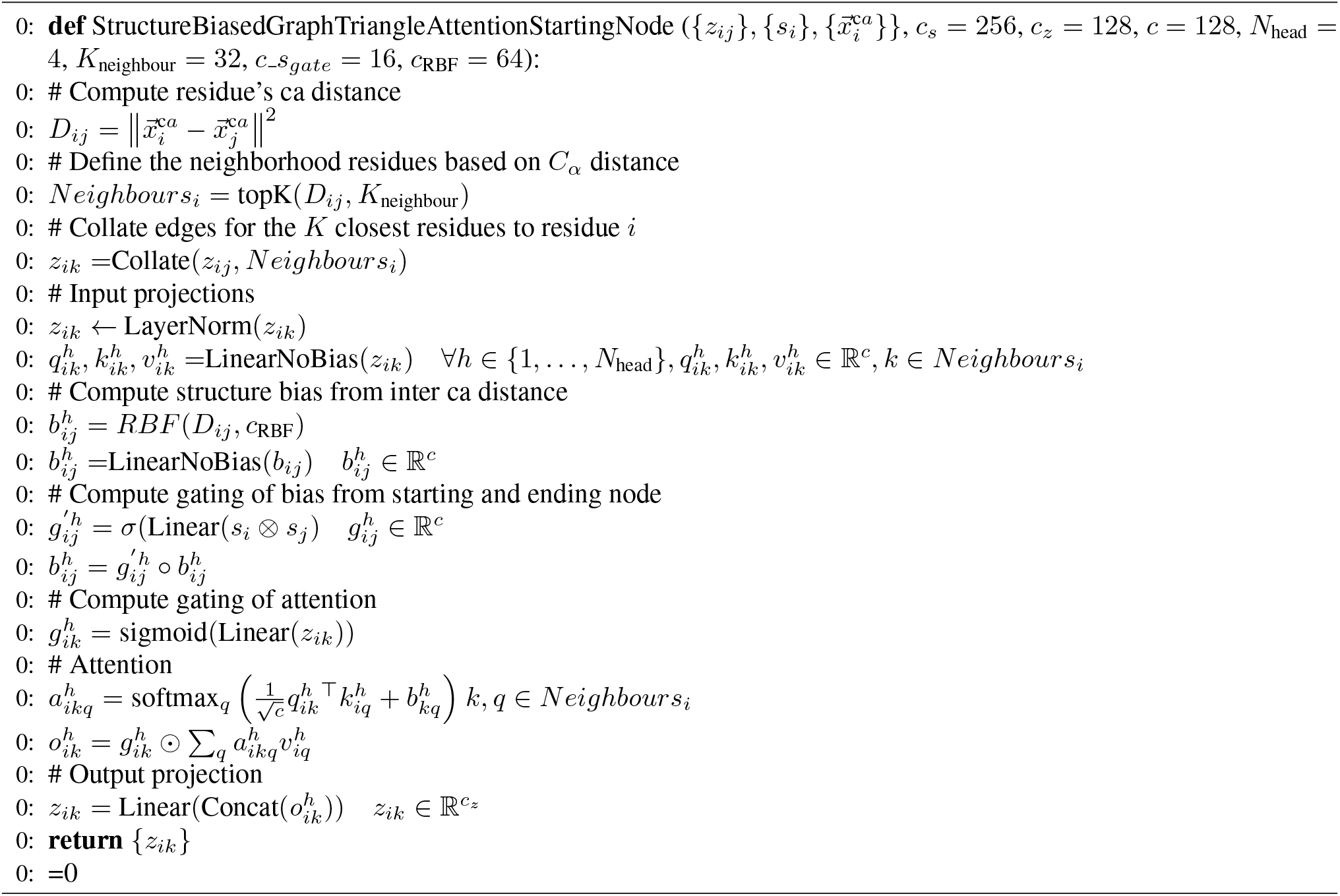

In the graph triangle block, the neighbor k residues of each residue are selected based on the inter C_*α*_ distance of current *T*_*ℓ*_, which means the attention graph is dynamically constructed in the network. By leveraging the characteristics of the diffusion model and dynamic refinement of the graph, triangle graph attention can avoid inaccurate graph construction in the early step, where the input structure is very noisy. At the same time, the efficiency of graph triangle attention calculation is maintained. When calculating the attention between *k*_*ik*_ and *q*_*iq*_, the structure bias is computed from 64 bins of radial basis function (RBF) equally spaced from 0Å to 32Å for distances between Ca for k and q residue.

#### Score prediction and reverse sampling schedule

We describe how the density score of rotation and translation of frames is calculated and how the SDE structure denoiser is applied to the sampling from noise mentioned in Figure 2 and Algorithm 1.

At each timestep between 0 and 1, after *L* layers we take the final frame *T*_*ℓ*_ as the predicted clean structure of *t* = 0

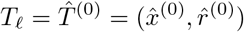

With the prediction 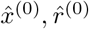 and the input frames *T* ^(*t*)^ = (*x*^(*t*)^, *r*^(*t*)^) of current timestep, ℝ ^3^ and SO(3) score of residue n is computed by

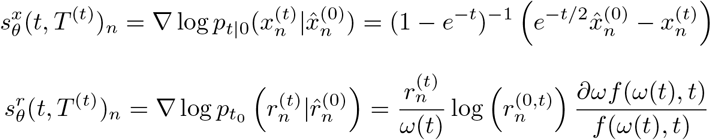

To define the formulation of the reverse process, we first set up the forward scheduling of by

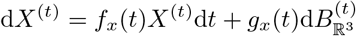

This equation describes the forward process of translation, where 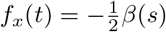 is drift coefficient and 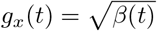 is diffusion coefficient. We choose linear schedule with introduced by Song et al. and Ho et al. β _min_ = 0.1, β _max_ = 20 as the SDE scheduler of translation.

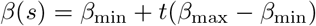

By defining this linear schedule. with 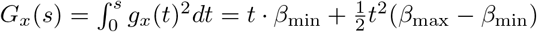, the distribution of translation at timestep t can be written as

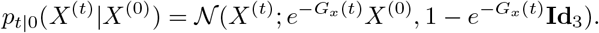

The forward diffusion process of rotation can be written similarly but without the drift coefficient term since it is in SO(3) space

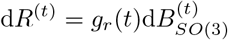

By introduce a time scaling factor 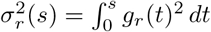 and 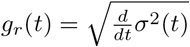, the probability distribution of rotation can be described as

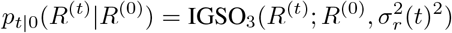

where

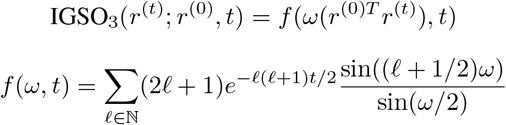

ω(*r*) is the rotation angle in radians for any *r* ∈ SO(3). By defining σ_*r*_(*t*) = log (*t* · exp {σ_max_} + (1 *− t*) exp {σ_min_}), we are able to control the rotation diffusion schedule through *g*_*r*_(*t*) with σ_min_ and σ_min_. We take the same setting 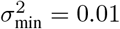 and 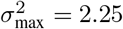 as described in FrameDiff.

With the forward process defined above, the SDE sampling procedure of translation and rotation can be described as

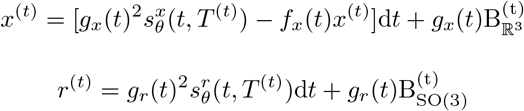

#### Noise scaling

To improve the backbone designability while keeping the diversity, we sample various noise levels in Table 3, noise level is applied to the Brownian motion of ℝ^3^ and SO(3) as

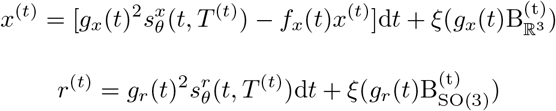

where noise level is noted as ξ *∈* [0, 1] in the Table 3 and equation above.

##### Algorithm 3

Structure biased graph triangle attention around ending node

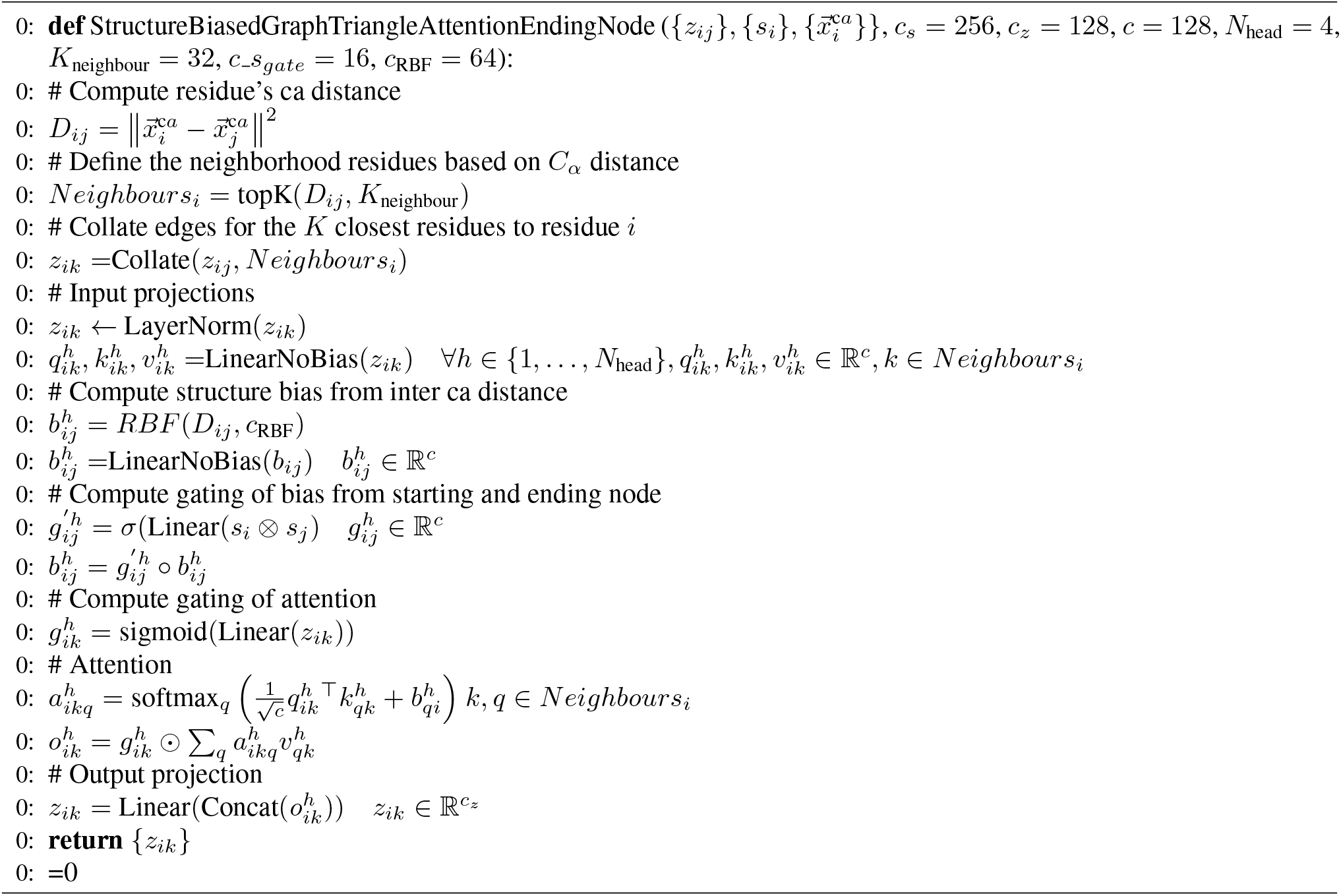

**Table 6:**
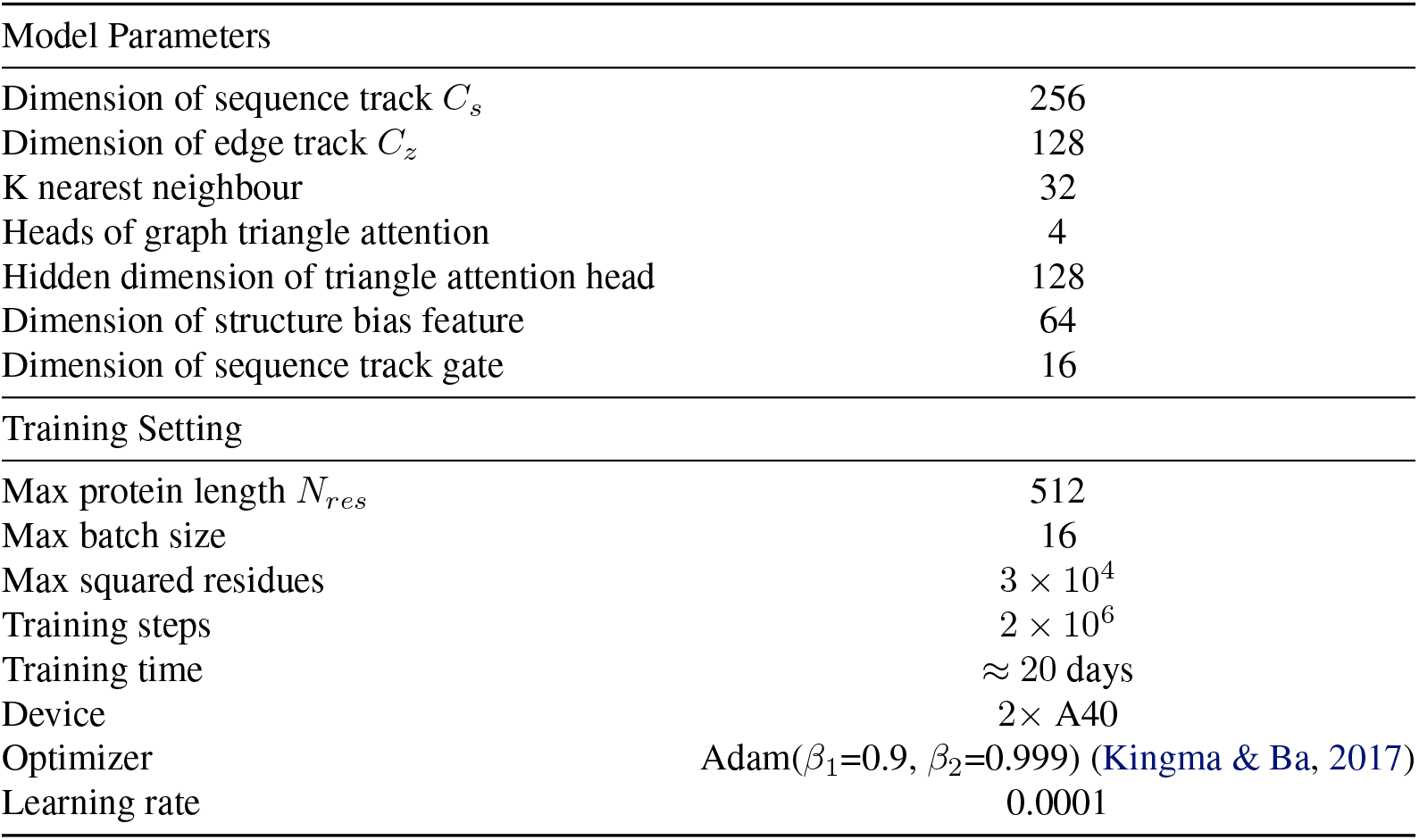
Model Training Parameters

## B. In vitro experiment protocol

### Protein purification and expression

Synthetic genes encoding designed protein were purchased from Universe Gene Technology. These sequences were cloned into the pET28a vector, included N-terminal or C-terminal histidine tags and an HRV 3C protease cleavage site. These plasmids were transformed into BL21 (DE3) E. coli competent cells. All transformants were cultured into 50 ml of LB medium with 50 mg/ml kanamycin. Protein expression was induced with 1 mM isopropyl 1-thio--d-galactopyranoside at 37 °C overnight or at 20 °C overnight after initial growth for 6 to 8 h at 37 °C. The cells were harvested by centrifugation and lysed by sonication after resuspension of the cells in lysis buffer (25 mM Tris pH 7.0, 150 mM NaCl). The cell lysate was cleared by centrifugation (12,000 × rpm). The supernatant was purified by 1 ml Ni2+ immobilized metal affinity chromatography with Ni-NTA Superflow resin (Qiagen). Resins with bound cell lysate were washed five times with 5 mL of washing buffer (comprising lysis buffer and 30 mM imidazole) and eluted with 6 mL of elution buffer (comprising lysis buffer and 300 mM imidazole). Both eluates were analyzed using 15% SDS-PAGE gel to assess purity. The histidine tags were cleaved using histidine-tagged HRV 3C protease during dialysis against lysis buffer overnight. A second IMAC purification was performed for HRV 3C cleaved samples to capture uncleaved protein and HRV 3C protease. Designs were finally purified using Superdex 200 Increase 10/300GL (GE Healthcare) with lysis buffer.

### Circular dichroism experiments

Circular dichroism spectra were recorded on a Chirascan V100 circular dichroism spectrometer (Applied Photophysics) using protein concentrations ranging from 0.6 to 0.9 mg/ml. Thermal melt analyses were conducted over a temperature range of 25°C to 95°C, measuring CD at 222 nm. Wavelength scans (190 to 260 nm) were recorded at both 25°C and 95°C. All reported measurements were obtained within the linear range of the instrument.

For crystallization, the plasmids were transformed into BL21 (DE3) E. coli competent cells. The transformants were cultured in 10 ml of LB medium with 50 mg/ml kanamycin at 37 °C overnight. The cultures were transferred to 1L of LB medium with 50 mg/ml kanamycin and incubated at 37 °C. Protein expression was induced with 1 mM IPTG at 37 °C overnight. Protein purification steps were carried out as described above.

### Crystallization, data collection, and structure determination

The crystals were grown using the hanging drop method at room temperature (18 °C). The drops consisted of 1 L of 40 mg/ml protein and 1 L of precipitant solution (100 mM Tris pH 8.5, 200 mM NaCl and 30For diffraction, the crystals were transferred into a solution containing 20% glycerol as a cryoprotectant. Subsequently, the crystals were loaded onto the X-ray diffractometer (Rigaku, XtaLAB Synergy Customer). The diffraction data was collected at 100 K and processed with the reduction program CrysAlisPro. The structures were solved by molecular replacement using Phaser in PHENIX8. The structures were manually refined with Coot9 and PHENIX10.

**Figure 5:**
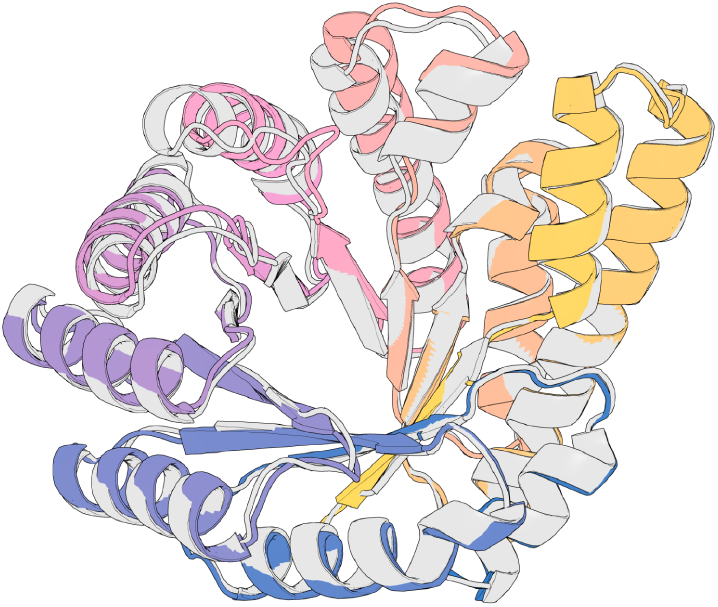
The alignment of experimentally solved crystal structure and diffusion model of 300aa_3, crystal backbone(grey) is overlaid with diffusion backbone(colors) with a global C_*α*_-RMSD 0.91 Å. The setting of data collection and structure determination is provided in Table 7

**Table 7:** Data collection and refinement statistics.

| Parameter | Value |
| --- | --- |
| Name | 300aa_3 |
| Wavelength | 1.541 |
| Resolution range | 29.13 - 1.8 (1.864 - 1.8) |
| Space group | P 1 21 1 |
| Unit cell | 46.4585 76.1478 83.3196 90 91.9037 90 |
| Total reflections | 247022 (15772) |
| Unique reflections | 53811 (4931) |
| Multiplicity | 4.6 (2.9) |
| Completeness (%) | 97.87 (91.26) |
| Mean I/sigma(I) | 24.16 (3.29) |
| Wilson B-factor | 16.89 |
| R-merge | 0.06467 (0.49) |
| R-meas | 0.07301 (0.5975) |
| R-pim | 0.03309 (0.338) |
| CC1/2 | 0.975 (0.488) |
| CC* | 0.994 (0.81) |
| Reflections used in refinement | 52709 (4928) |
| Reflections used for R-free | 1368 (129) |
| R-work | 0.1827 (0.2144) |
| R-free | 0.2163 (0.2486) |
| CC(work) | 0.961 (0.892) |
| CC(free) | 0.936 (0.880) |
| Number of non-hydrogen atoms | 5174<br>macromolecules: 4418<br>ligands: 0<br>solvent: 756 |
| Protein residues | 600 |
| RMS(bonds) | 0.007 |
| RMS(angles) | 0.90 |
| Ramachandran favored (%) | 99.33 |
| Ramachandran allowed (%) | 0.50 |
| Ramachandran outliers (%) | 0.17 |
| Rotamer outliers (%) | 0.64 |
| Clashscore | 7.73 |
| Average B-factor | 23.04<br>macromolecules: 21.76<br>solvent: 30.55 |

**Figure 6:**
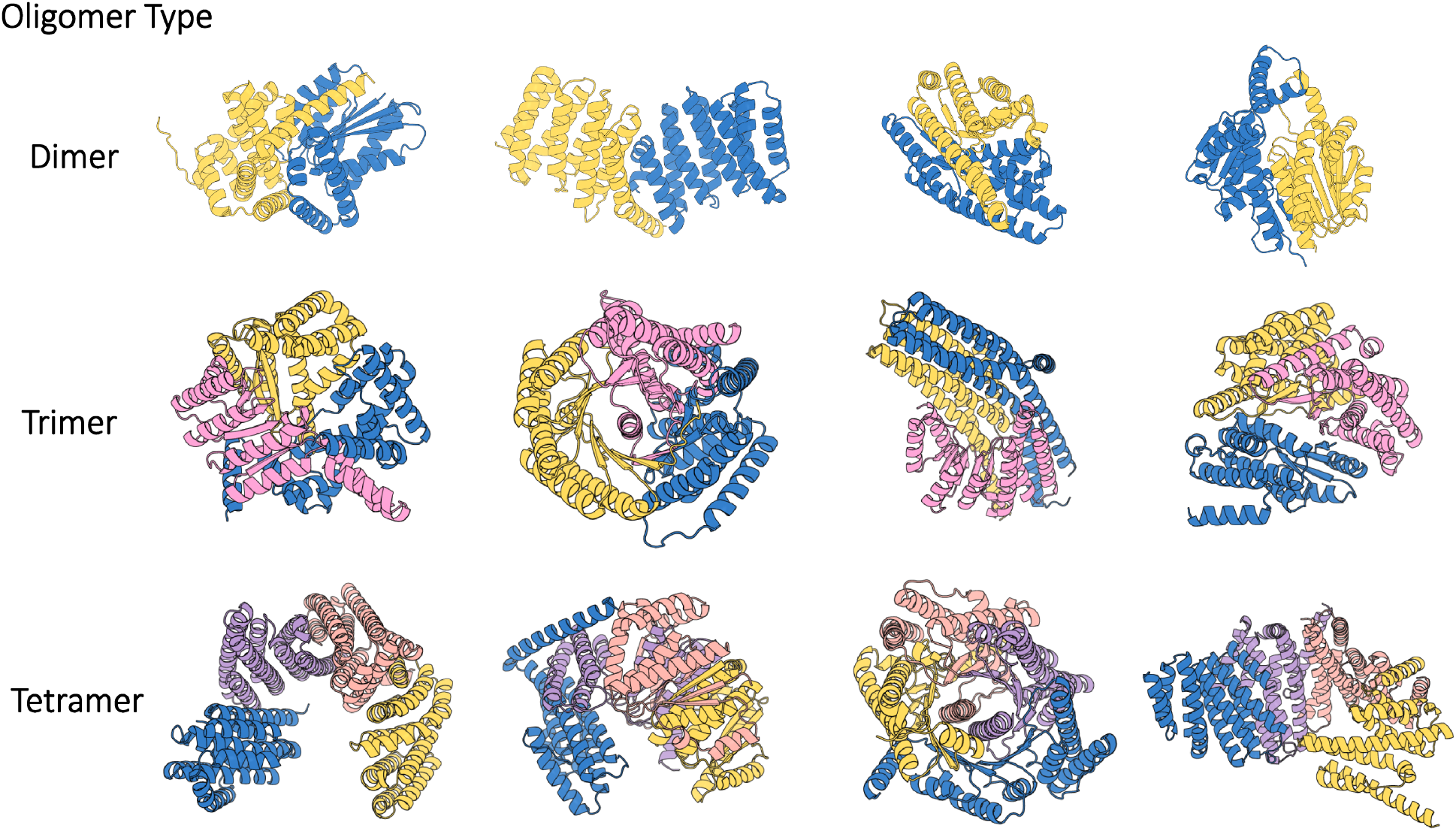
Visualization of oligomer samples across dimer, trimer, and tetramer. each chain is fixed at 200 residues with different colors, as shown in the figure.

